# Regional and Temporal Changes in Early Structural Remodeling Following Myocardial Infarction via Semi-Automatic Image Analysis

**DOI:** 10.1101/2025.08.23.668999

**Authors:** Catherine C. Eberman, Yuming Liu, Kevin W. Eliceiri, Colleen M. Witzenburg

## Abstract

Reperfusion therapy, the restoration of blood flow following a myocardial infarction (MI), is one of the most effective treatment strategies. Unlike early reperfusion therapy, differences in infarct size or collagen content have not been reported in late reperfusion therapy. To evaluate the spatial-temporal effects of late reperfusion therapy, we conducted multimodal imaging of histologic sections of rat myocardium following permanent coronary artery occlusion or three hours of occlusion. Semi-automatic partitioning identified the infarct core, infarct border, and healthy periphery regions from label-free liquid crystal based polarized light microscopy (PolScope) images taken throughout the first 5 days of healing. Associated brightfield and standard polarized light microscopy images of hematoxylin-eosin or Picrosirius Red stained sections were also used to determine cellular and collagen fiber densities, respectively. Even when we consider multiple definitions for the vulnerable infarct border, its size decreased faster in late reperfusion therapy samples. Temporal patterns in collagen density also indicated late reperfusion led to a more rapid progression through the necrotic phase of healing (when the infarct is vulnerable to rupture) and earlier progression to the fibrotic phase of healing (when the infarct stabilizes). Notably, we also observed a broader region of provisional non-collagenous matrix in late reperfusion samples during the necrotic phase of healing. Together these findings suggest late reperfusion therapy accelerates healing and potentially changes the spatial pattern of provisional matrix deposition during the period the heart is most susceptible to rupture events.

## 1. Introduction

Each year over 800,000 people in the United States have a heart attack or myocardial infarction (MI) which involves occlusion of coronary artery blood flow, myocardial ischemia, and death of the downstream cardiac muscle [1]. After coronary artery occlusion, the affected myocardium undergoes a robust inflammatory response that ultimately leads to significant changes in tissue content and structure. This highly dynamic response is frequently separated into three healing phases: the inflammatory or necrotic phase, the proliferative or fibrotic phase, and the maturation or long-term remodeling phase [1], [2], [3]. First, as cardiomyocytes become necrotic, they release damage-associated molecular patterns and pro-inflammatory cytokines that attract neutrophils and macrophages to the infarction site. These infiltrating immune cells then secrete matrix metalloproteinases (MMPs), which degrade the surrounding extracellular matrix (ECM), enabling them to degrade and remove necrotic cardiomyocytes[4]. The robust native ECM is initially replaced with a weak provisional matrix [2], [5]. A marked rise in fibroblast infiltration and collagen deposition signifies the transition to the fibrotic phase of healing.

While the post-MI inflammatory response is critical to healing, it renders the infarcted myocardium vulnerable to risk of cardiac rupture [6], [7], [8], [9].Reperfusion therapy, the restoration of blood flow, is one of the most effective treatment strategies for MI. Widespread use of reperfusion therapy has been corelated with a dramatic decrease in cardiac rupture [10], [11], [12], [13], [14], [15], [16] even when administered late, hours following MI and after cardiomyocyte death [14], [17], [18], [19], [20], [21], [22], [23]. Late reperfusion therapy does not reduce infarct size or transmurality [24], [25], [26], [27], [28]. Increased MMPs secretion and lower collagen content has widely been proposed as a potential rupture predictor [29], [30], [31], [32], but detailed studies on rodents do not support it [33], [34], [35], [36]. Thus, the underlying mechanisms of late reperfusion’s success in reducing rupture incidence are not well understood [14], [37], [38], [39]. Unfortunately, while the mortality and complication rate of MI have dropped remarkably in the reperfusion era, hospital fatality rates following rupture have remained stagnant [40]. We hypothesize that late reperfusion therapy leads to smoother regional changes in the tissue content and structure following an MI [41], [42], [43] increasing the loading threshold for a tear to form and propagate through the tissue.

Furthermore, spatiotemporal heterogeneity in infarct healing is an active area of investigation, particularly for therapies aimed at reducing long term heart failure risk [3], [44], [45].Following an MI significant regional heterogeneity is present, however, there has been substantial debate as to what constitutes the infarct core, border, and peripheral regions [46], [47]. In this study, we quantified the temporal changes in regional heterogeneity following permanent coronary occlusion and late reperfusion therapy (temporary coronary occlusion) in rats. We developed a semi-automated image processing approach to quantify differences in both cellular and collagen density in the longitudinal-circumferential plane of the ventricular wall during the first 5 days following an MI. Additionally, we investigated the impact of conservative and liberal variations in the definition of the infarct core, border, and peripheral regions on this spatial distribution of these measurements. Therefore, we were able to identify both temporal and regional differences throughout the healing process between the permanent occlusion and late reperfusion groups.

## 2. Methods

### 2.1 Animal Experiments

For this study, 31 male Sprague-Dawley rats (375-575g) were subject to infarction by ligation of the left anterior descending coronary artery. The artery was permanently occluded (PO) in 15 animals. In the other 16 animals the ligation was removed 3 hours post-occlusion to simulate late reperfusion therapy (LRT). All studies were performed in accordance with the Guide for the Care and Use of Laboratory Animals and reviewed and approved by the University of Wisconsin Institutional Animal Care and Use Committee. Hearts were harvested on day 1, 3, and 5 post-infarction. Each heart was arrested by retrograde aortic perfusion with phosphate-buffered saline and frozen at -80 degrees. Hearts from five animals, four from the LRT group and one from the PO group, were excluded from further analysis due to an insufficient infarct (less than 2mm^2^/g LV) or the presence of a substantial hematoma.

### 2.2 Dissection and Histology

Upon thawing, the left ventricle (LV) was dissected from the heart. The LV was then cut open longitudinally along the septum and the LV free wall was flattened by relief cuts at the apical edge (Figure S1). The LV was fixed in 10% formalin for 24 hrs and transferred to 70% ethanol briefly prior to histological staining. Histology sections, 5 μm in thickness, were cut in the longitudinal-circumferential plane of the mid-wall. Adjacent sections were stained with either hematoxylin-eosin (H&E) for general morphology or Picrosirius Red (PSR) for collagen fibers.

### 2.3 Multimodal Imaging

Images of the H&E stained slides were acquired using an upright microscope (Nikon Eclipse Ni-U) in brightfield mode with a 4x objective (Nikon Plan Flour 4x, NA=0.13) and Aperio AT2 Digital Pathology Scanner (Leica Biosystems) with a 20x objective (Plan Apo 20x, NA=0.75) to determine cellular density. Standard polarized light microscopy images of the PSR stained slides were acquired using the upright microscope in polarization mode with a 10x objective (Nikon Plan Flour 10x, NA=0.30) to determine collagen density. Label free liquid crystal based polarized light microscopy (PolScope) images of the H&E stained slides were acquired using a customized PolScope system in birefringence mode with a 4x objective (Nikon CFI Achro P 4x, NA=0.1) to assign infarct core, infarct border, and healthy peripheral regions. Additional regions of interest acquired with a 20x objective (Nikon CFI Achro LWD 20x, NA=0.4).

The configuration of the PolScope system was described previously [48]. Briefly, the PolScope system was built around a Nikon Eclipse TE200 with the addition of a liquid crystal (LC) based polarization state generator and a circular polarization state analyzer. Compared to the standard polarized light microscope, PolScope can generate any polarization state by the electrically controlled LC-based polarizer and model selected polarization states to computationally detect birefringent structures (such as collagen fibers) orientated at different directions with higher sensitivity from either unstained tissues or H&E stained tissues. While collagen fibers could be identified in PolScope images in the presence of muscle cells these fibers could not be easily isolated (Figure 1A). Standard polarized imaging of PSR stained slides enabled identification of collagen fibers throughout the tissue regardless of the surrounding muscle content (Figure 1B). Additionally, the current version of PolScope in birefringence mode is not generally used for PSR stained samples since this stain introduces dichroism in collagen fibers that would be calculated as birefringence in PolScope.

**Figure 1:**
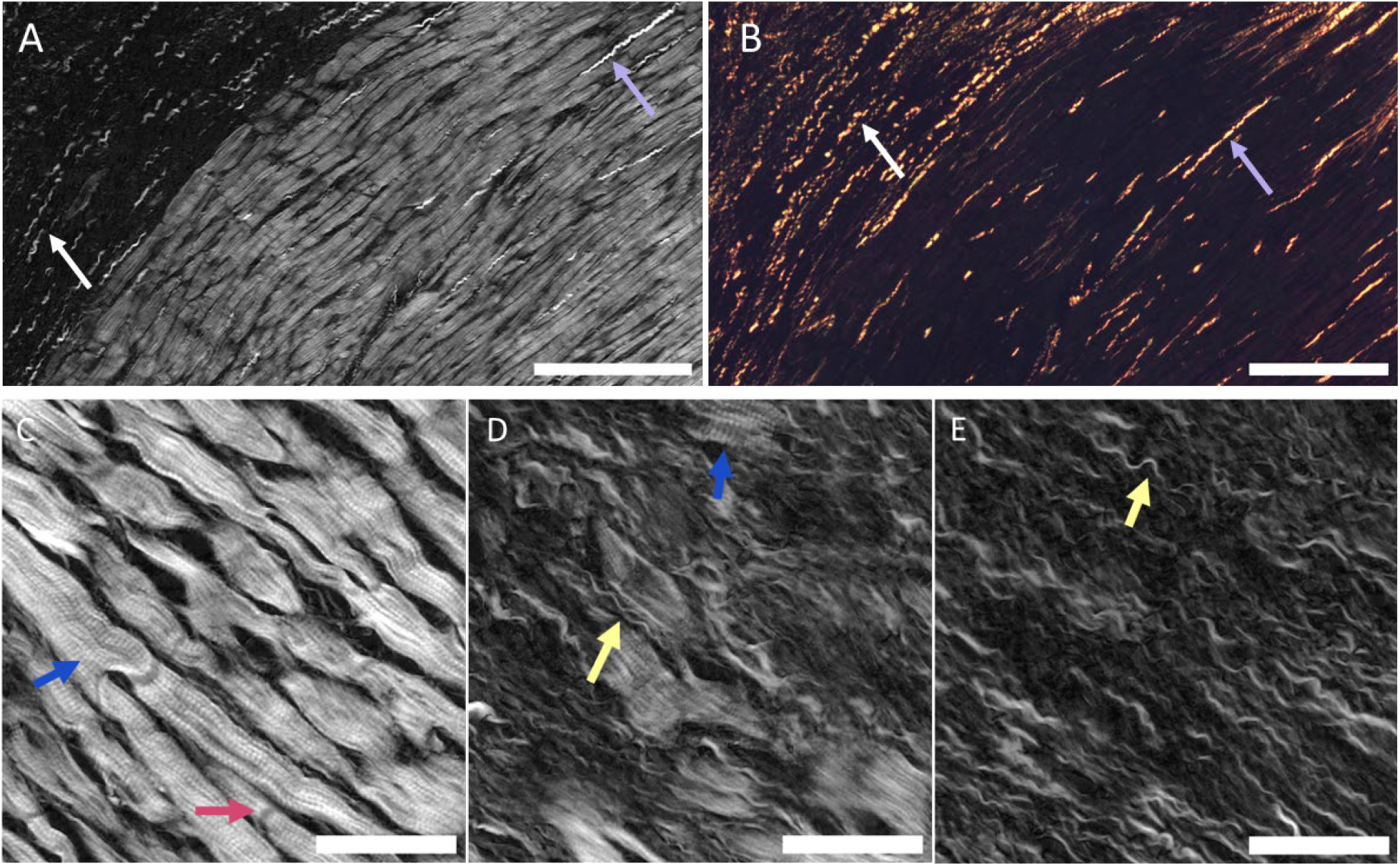
Collagen fibers within a representative PolScope image of an H&E stained section (A) and the standard polarized light microscopy image of the neighboring PSR stained section (B). White arrows identify collagen fibers in an area with muscle loss and purple arrows identify collagen fibers in regions with muscle. PolScope allows for higher quality imaging of collagen fibers within the core, however in regions with healthy cardiomyocytes they are obscured. A, B Scale bar = 250μm. PolScope provides discriminant features among peripheral region (C), border region (D) and core region (E) of the left ventricle (LV). The tissue section was from a representative rat five days following permanent left anterior descending coronary artery occlusion. The peripheral region contained myocytes that appear healthy with visible sarcomere units (blue) and intercalated discs (red). The border region contained damaged myocytes with irregular shapes and less distinct features as well as an elevated amount of collagen fibers (yellow). The core region contained no myocytes and had an abundance of collagen. C, D, E Scale bar = 50μm

The PolScope system provided superior visualization of cardiomyocytes (Figure 1C), enabling identification of individual sarcomere units and intercalated discs within bright regions of healthy cells. Damaged regions were darker and less defined (Figure 1D), and the infarct core was black with no visible cardiomyocytes (Figure 1E). Therefore, regional topology was determined from PolScope images of H&E stained sections.

### 2.4 Image Processing and Analysis

Figure 2 outlines the image processing and analysis procedure utilized to quantify regional collagen and cellular area fractions throughout the infarcted samples from PO and LRT groups at days 1, 3, and 5 post-infarction.

**Figure 2:**
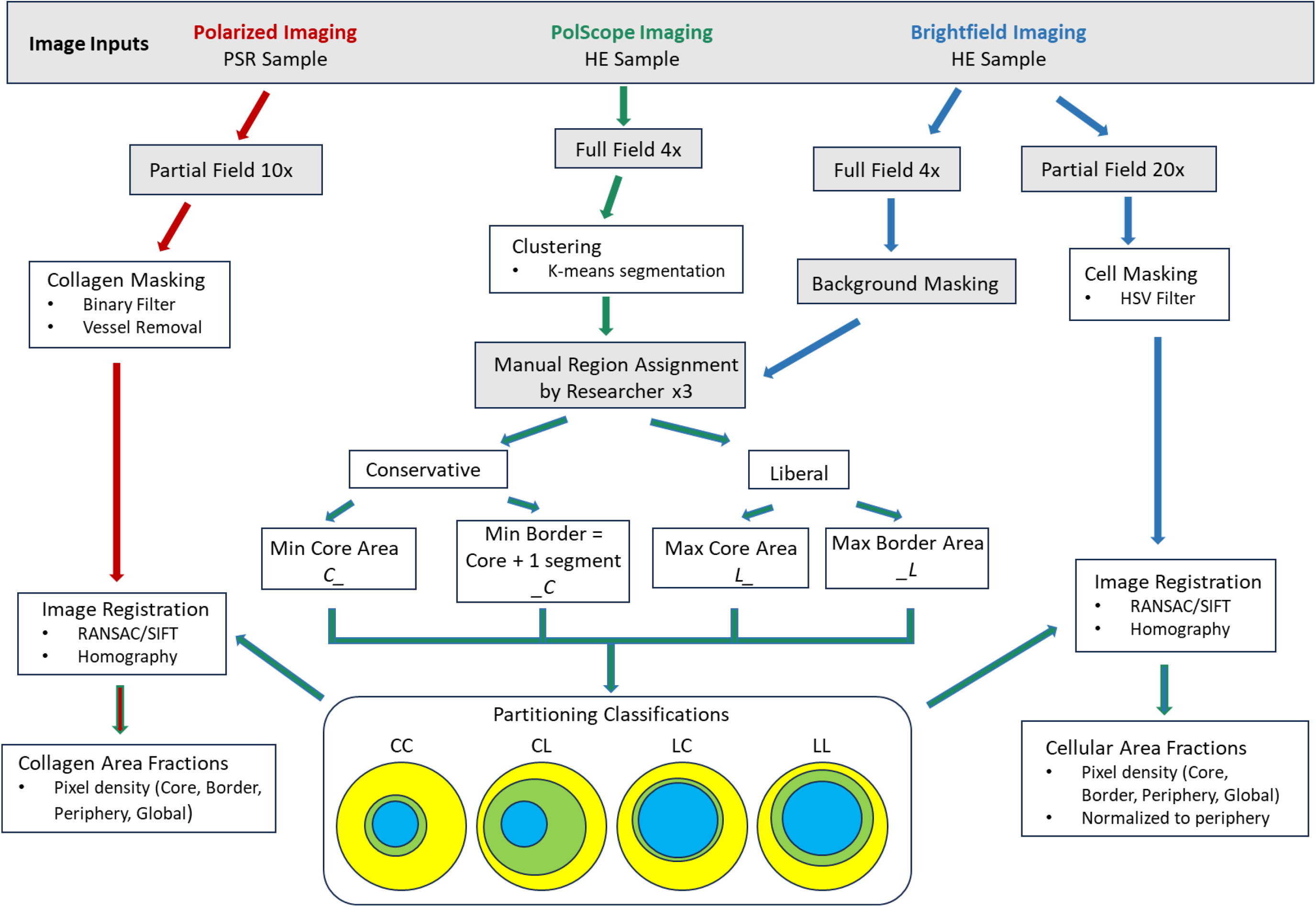
Schematic outlining the image processing and analysis procedure utilized to quantify regional collagen and cellular area fractions. PolScope images underwent automated clustering to determine 15 separate areas with similar intensity. Next, three researchers, blinded to day and group, assigned each cluster to either infarct core, infarct border, or healthy periphery. Disagreement between these definitions lead to four partitioning classifications which spanned the most conservative and liberal core and border interpretations (C_ and _C indicate conservative segmentations, defined as the minimum core area chosen by any researcher and one segment larger than the given core area. L_ and _L indicate liberal segmentations defined as the maximum core and maximum border areas chosen by any of the researchers). Partitioned PolScope images were then registered to the appropriate brightfield image and regions were mapped to identify cellular and collagen densities. Image processing procedures within grey boxes involved manual input while those shown in white boxes were fully automated.

#### 2.4.1 Cell Masking

Cell infiltration was quantified from brightfield imaging (Aperio scanner) of H&E stained sections [49]. Color-based thresholding in the HSV color space identified and isolated cell nuclei [50]. Pixels were included as ‘cells’ if the following was true:

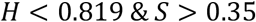

where *H* indicates hue and *S* indicates saturation. Visual inspection indicated cell nuclei were circular in shape and approximately 60 to 100 pixels in area. Therefore, we automatically detected and removed regions less than 8 pixels in area that were the same color as cells. Regions with an eccentricity of greater than 0.9, which were often associated with bands of muscle, were also removed. Lastly, any small holes within cell nuclei were filled in.

#### 2.4.2 Collagen Masking

Collagen density was quantified from standard polarized light microscopy images of PSR stained sections. The PSR stain binds specifically to collagen fibrils within myocardial tissue [51], [52]. Under linearly polarized light, the birefringence of the muscle is low compared to the collagen fibers. To determine collagen density a binarization was applied to the PSR images. Briefly, images were converted to grayscale, and a binarization threshold of 0.3 was applied, as this reliably removed the background coloration without elimination of collagen fibers. Next, vessels and image edges, which exhibited high collagen fiber concentrations, were removed to avoid skewing local densities. To identify vessels and image edges, we applied a median filter to smooth the original PSR image followed by a binarization filter. All objects smaller than 20,000 pixels were removed, leaving a mask containing only the large dense vessels and sample edges. This mask was then subtracted from the binarized PSR image to produce a final binarized image containing only the myocardial collagen.

In PSR stained images, color indicates fiber type, with collagen I appearing redder and collagen III greener [52]. To analyze color variation within PSR stained samples, the images were converted to the LAB color scale. The background was masked out using a binarization filter, and average color was determined for the A axis, which indicates the green-to-red color transition.

#### 2.4.3 Clustering and Background Masking

An intensity-based approach was developed to identify areas of each sample within the infarct core, infarct border, and healthy periphery. PolScope images were processed with a median filter to create a blurred version [53]. The blurred image and the original image were then inputted into the k-means segmentation algorithm in MATLAB [50] to produce 15 intensity-based clusters. We selected 15 clusters because this number was sufficient to isolate regions of moderate intensity while not overly segmenting regions of interest. The algorithm created clusters such that variance was minimized within the given cluster,

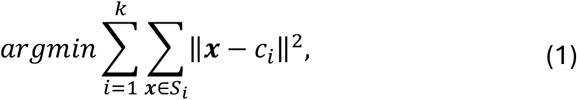

where k was the number of clusters, 15, **x**, was the points within a cluster, S was the set defining the cluster, and *c*_*i*_was the centroid of cluster *i* [53]. Clusters were then sorted by intensity level, with the darkest given a value of 1 and the brightest a value of 15. Following sorting, the cluster matrix was run through a smoothing filter to reduce the effects of individual pixel variation on the final masks. A background mask was created using H&E brightfield imaging to identify and remove image regions lacking tissue such as borders and vessel gaps.

#### 2.4.4 Manual Region Assignment and Partitioning Classifications

For each sample, three independent researchers, who were blinded to sample group or day, manually assigned each of the 15 clusters to either the infarct core, infarct border, or healthy peripheral region. Consistent with the ongoing debate about what constitutes the border zone around an infarct [47], [54], there were wide discrepancies and high interobserver variability. Therefore, to evaluate the bounds of what constitutes the infarct core and boundary we utilized four different partitioning classifications consisting of a conservative or liberal definition of the infarct core and border (C and L in Figure 2, respectively where the core definition is first and the border second). Conservative core regions (C _ in Figure 2) were defined to include clusters that were distinctly core only and did not contain any cardiomyocytes. In contrast, liberal core regions (L _) were defined to include all clusters containing core and therefore may have contained some damaged or degraded cardiomyocytes. Similarly, the liberal border region was defined as the largest region that contained any signs of cardiomyocyte irregularity (_ L). Whereas the conservative border region was defined as one incrementally brighter layer beyond the core region (_ C).

Border size was quantified following region assignment. The distance of every border point to the nearest non-border point was determined via a MATLAB algorithm for Euclidean distance [55]. Then, the local maximum distance across the border zone was identified to find the farthest points, which represent the center of the region. The average of these maximum distances was doubled to determine the width of the border.

#### 2.4.5 Image Registration and Regional Mapping

To determine core, border, and peripheral collagen and cellular density within the LV samples the regional masks associated with each partitioning classification were mapped to the standard polarized light PSR and brightfield H&E images. To register these image pairs shared points were determined. When mapping PolScope regions to H&E images automated feature recognition was accomplished using SIFT [56], [57] and point selection was accomplished using RANSAC [58]. To run SIFT on the H&E and PolScope images they were first binarized to increase feature similarity using a threshold of 0.05 for PolScope and 0.65 for H&E. Images were then downsized by a factor of 4 (PolScope) or 10 (H&E) to reduce runtime. When mapping PolScope regions to PSR images, however, the higher degree of feature variation required manual identification of shared points. Next, a homography matrix, *M*, was calculated from the shared points [59] and applied to the PolScope derived topology masks, *I*^*pol*^, transforming them to match either the H&E or PSR images. Additionally, a transformation, *T*, was applied to *M* return image coordinate system to full scale (Figure 3).

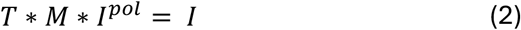

**Figure 3:**
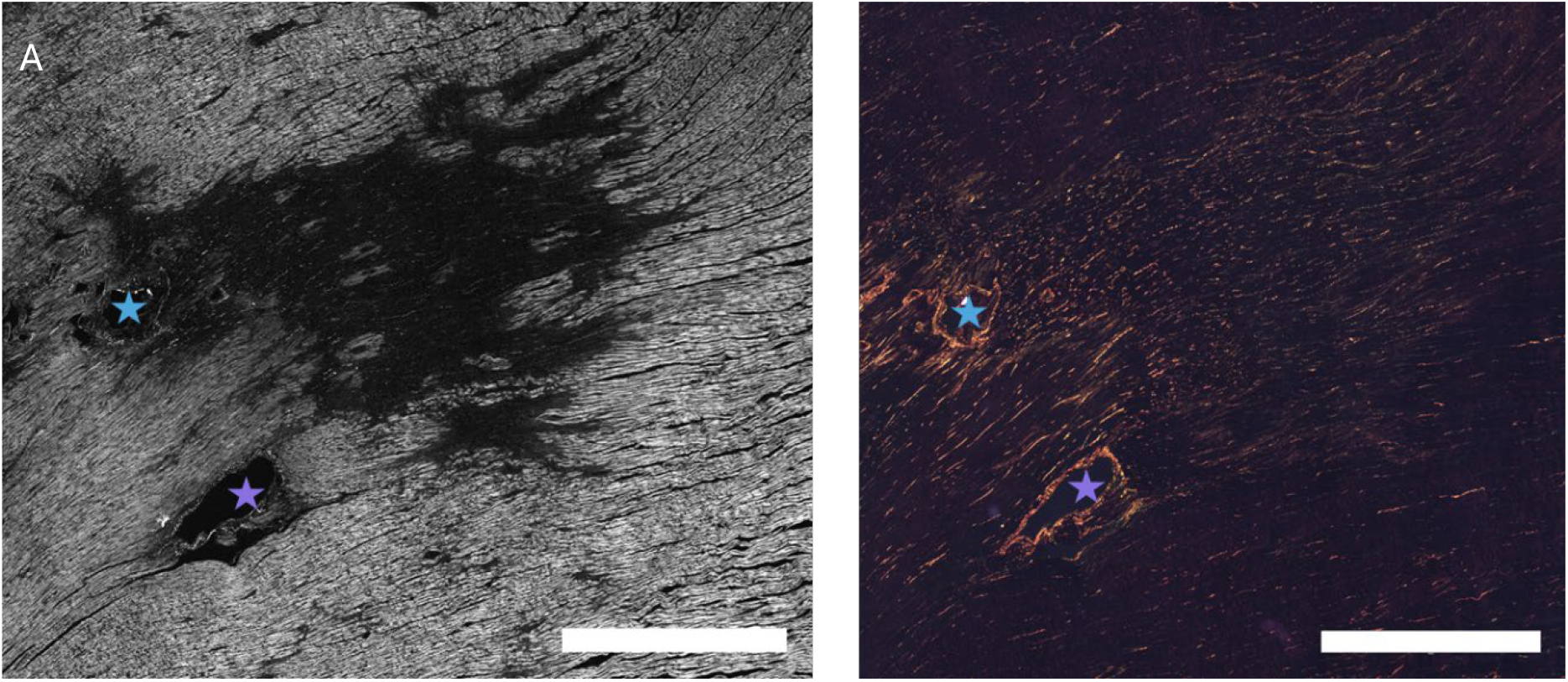
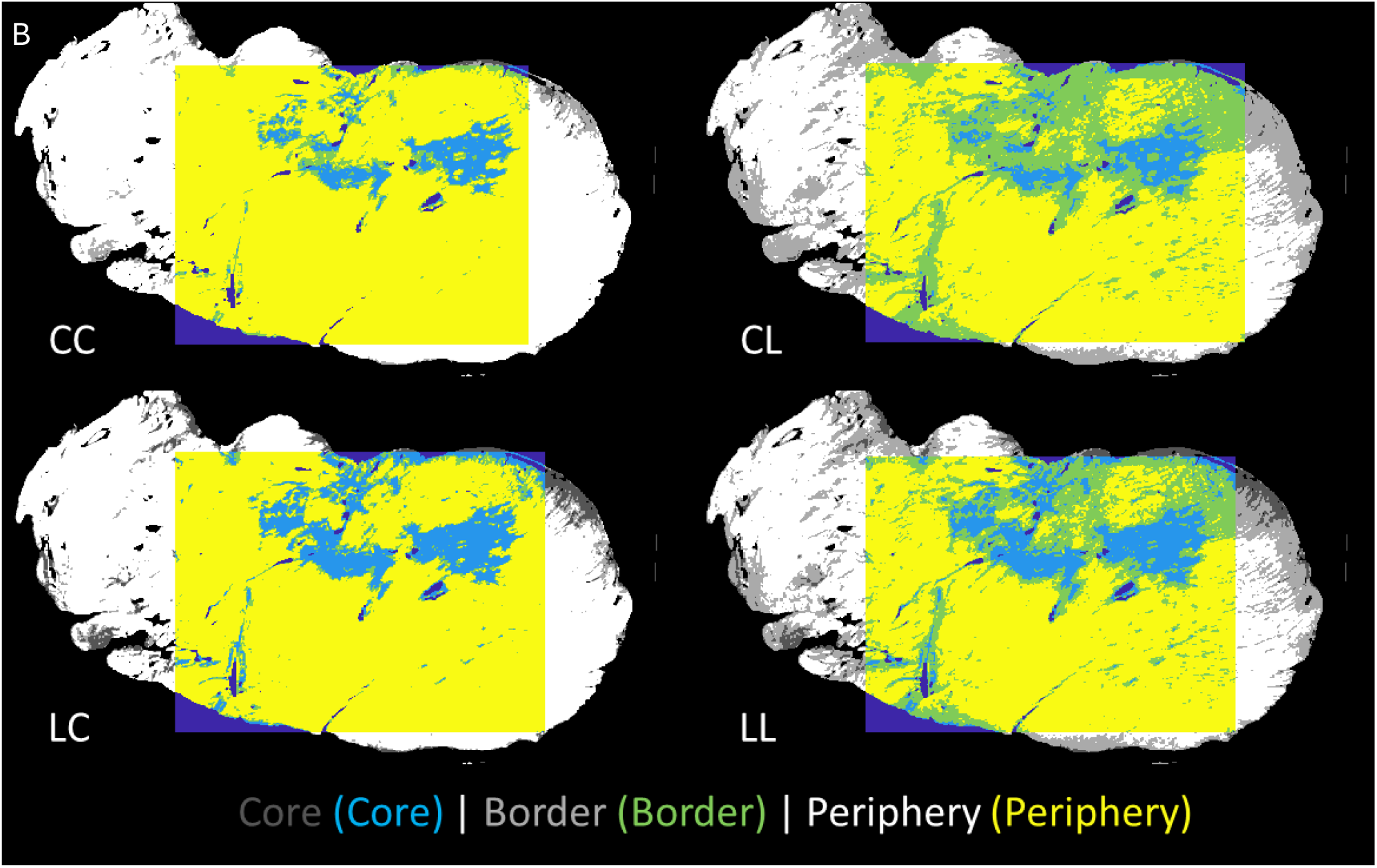
A) Comparison of a representative PolScope image of an H&E stained section (left) and a standard polarized light microscopy image of the neighboring PSR stained section (right) containing an infarct. Stars identify shared landmarks within the slices. Muscle loss in core region is clearly visible via PolScope while standard polarized light imaging does not differentiate between these regions. Scale bar = 1500μm. B) The four different regional assignments for a representative PolScope image of an H&E stained section of myocardium from a late reperfusion animal 5 days post-MI. The top panels show the conservative segmentation of the infarct core (light blue) and the bottom show liberal segmentation of the infarct core. Similarly, the left shows the conservative segmentation of the border (green), and the right panels show the liberal border segmentation. The peripheral tissue is yellow, and dark blue indicates the image background. Colored regions indicate the analysis window. Partitioning classifications: CC – conservative core and border, CL – conservative core and liberal border, LC – liberal core and conservative border, and LL – liberal core and border.

Each transformed H&E or PSR image specific topology mask, *I*, was then applied to the appropriate cellular and collagen density results. Regional density was calculated for the infarct core, infarct border, and healthy periphery as well as a global density. Cellular density values were then normalized to the periphery to account for variations in segmentation thresholding.

### 2.5 Statistical Analysis

Statistical analysis was performed in RStudio 2024.12.1 and GraphPad Prism, version 10.1.2. To compare border width, a 2-way ANOVA with group (i.e. PO or LRT) and time point (1, 3 or 5 days post-MI) was conducted for each partitioning classification. A 3-way ANOVA was conducted for both cell and collagen density. Based on significant effects and interactions, subsequent 2-way ANOVAs were conducted for both cell and collagen density, with group and time point, time point and region (core, border, or periphery), and group and region for each partitioning classification. For collagen color and type within the infarct core, a 2-way ANOVA with group and time point was conducted for the conservative and liberal partitioning classifications. Post hoc multiple comparison testing was completed via Tukey testing where p-values less than 0.05 were considered significant, and p-values between 0.05 and 0.10 were considered trending towards significant. Summary statistics are reported as mean ± standard deviation.

## 3 Results

### 3.1 PolScope and Standard Polarized Light Microscopy Imaging

Figure 3A shows the infarct region of a representative H&E stained section of myocardium from a LRT animal 5 days post-MI imaged via PolScope as well as the neighboring PSR stained section imaged via standard polarized imaging. PolScope allows for visualization of the bright muscle fibers and enables clear delineation of the regions where the muscle has been damaged or removed. Therefore, the core, border, and peripheral regions of the infarct can be identified.

### 3.2 Semi-Automated Region Assignment

Figure 3B shows the four different regional assignments determined following automated intensity-based clustering and manual region assignment for the representative PolScope image of the H&E stained shown in Figure 3A. Due to the lack of formal definitions for the border region, both conservative and liberal interpretations were investigated. As anticipated, the combination of both conservative core and border definitions produced the smallest infarct-to-total tissue ratio (upper left panel) and the liberal core and border definitions produced the largest infarct-to-total tissue ratio (lower right panel) with other combinations in between (Table S1). Note that in these computations, the total tissue area was considered. However, when calculating border width as well as regional cell and collagen densities, only pixels within the analysis window was considered. The orientation of the muscle cells at the sample edge produced a different polarization intensity. Thus, healthy edge cardiomyocytes were often at a lower brightness than those in the sample bulk. To prevent the inaccurately including healthy edge cardiomyocytes in the infarct core or border regions and to reduce computational load, an analysis window (Figure 3B) was applied to minimize this effect while still capturing as much of the true infarct region as possible.

### 3.3 Border Width and Infarct Area

Border width, defined by maximum Euclidean distance, varied with partitioning classification as expected. In general, border width was smallest when the LC partitioning classification was applied and largest when the CL partitioning classification was applied (Figure 4).

**Figure 4:**
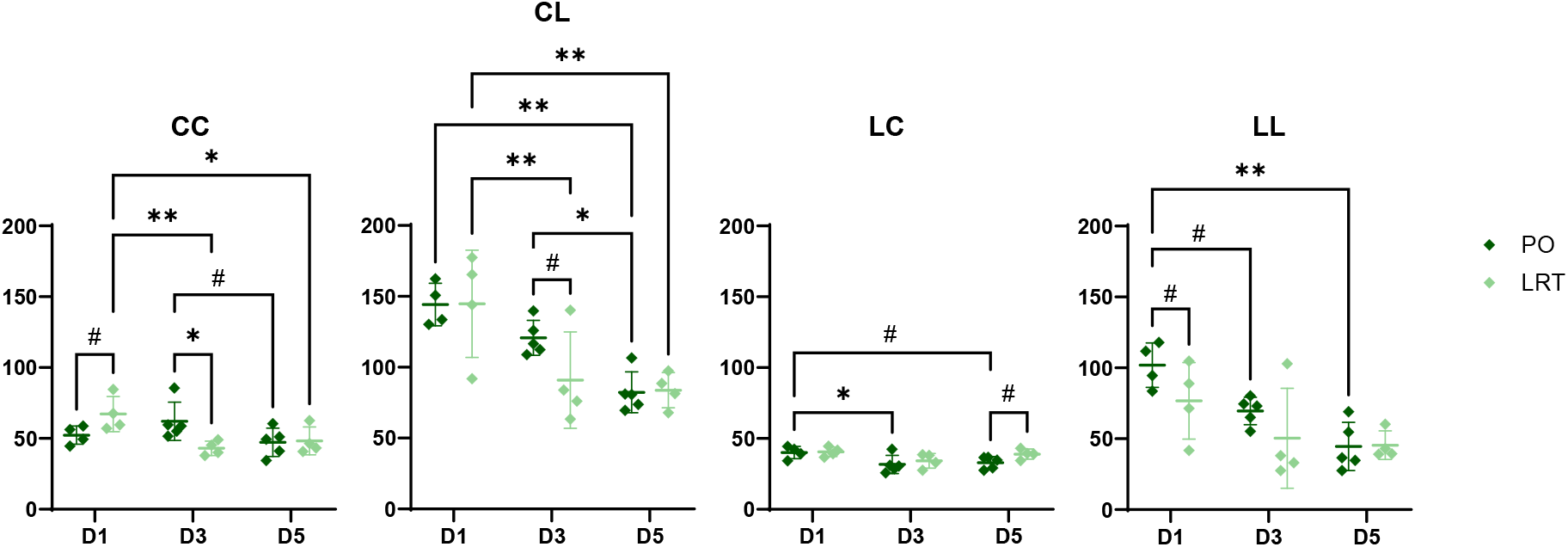
Border width (μm) for all partitioning classifications at all time points (days (D) 1, 3, and 5). There was a downward trend in border width (μm) over time and width decrease earlier in late reperfusion therapy (LRT) samples than permanent occlusion (PO) samples. Statistical significance is indicated for a 2-way ANOVA when the post hoc Tukey test indicated a pairwise comparison with # p < 0.1, ^*^ p < 0.05, ^**^ p < 0.01. The LRT group included 4 hearts per day, and the PO group included 4 hearts at day 1 and 5 hearts at days 3 and 5.

At day 1, paired comparisons from the 2-way ANOVA via Tukey tests indicated a trend towards a wider border in the LRT group when the conservative core and border (CC) partitioning classification was applied (p = 0.051), however this trend reversed when the liberal core and border (LL) partitioning classification was applied (p = 0.099). At day 3 for all partitioning classifications, the border was wider in the PO group than the LRT group, except LC where the border was similar in size (31.7 ± 6.4 μm and 34.3 ± 5.2 μm, respectively). Paired comparisons indicated the border was significantly wider in the PO group at day 3 for the CC classification (p = 0.011) and trended towards significance for the CL classification (p=0.064).

Paired comparisons from the 2-way ANOVA via Tukey tests also indicated temporal trends in border width. When the more liberal border definition was applied, width decreased with time and this downward trend progressed more quickly in the LRT group than the PO group. For the LRT group the border was significantly wider on day 1 than days 3 or 5 (p = 0.0086, 0.0032, respectively) whereas for the PO group it was significantly thinner on day 5 than days 1 or 3 (p = 0.0017, 0.030, respectively).

Core, border and total infarct areas were computed for each partitioning classification and there were no significant differences via 2-way ANOVA for any time point, or group. The only exceptions with subsequent Tukey testing were on day 5 for the liberal core area definition where the LRT group exhibited a larger core area than the PO group and on day 5 for the conservative core and border definition where the LRT group exhibited a significantly larger total infarct area than the PO group (Figure S2).

### 3.4 Cellular Area Fractions

Color-based thresholding reliably isolated cell nuclei from each image (Figure 5A and Figure S3). Regional area fractions were computed and normalized to the periphery for the analysis window for all four partitioning classifications. A statistical analysis via 3-way ANOVA indicated no significant interaction between group, time point, and region, but did indicate a significant interaction between time point and region for the CL and LL partitioning classifications (p= 0.0055 and p= 0.027 respectively). The 3-way ANOVA showed no significant difference with group for any classification.

**Figure 5:**
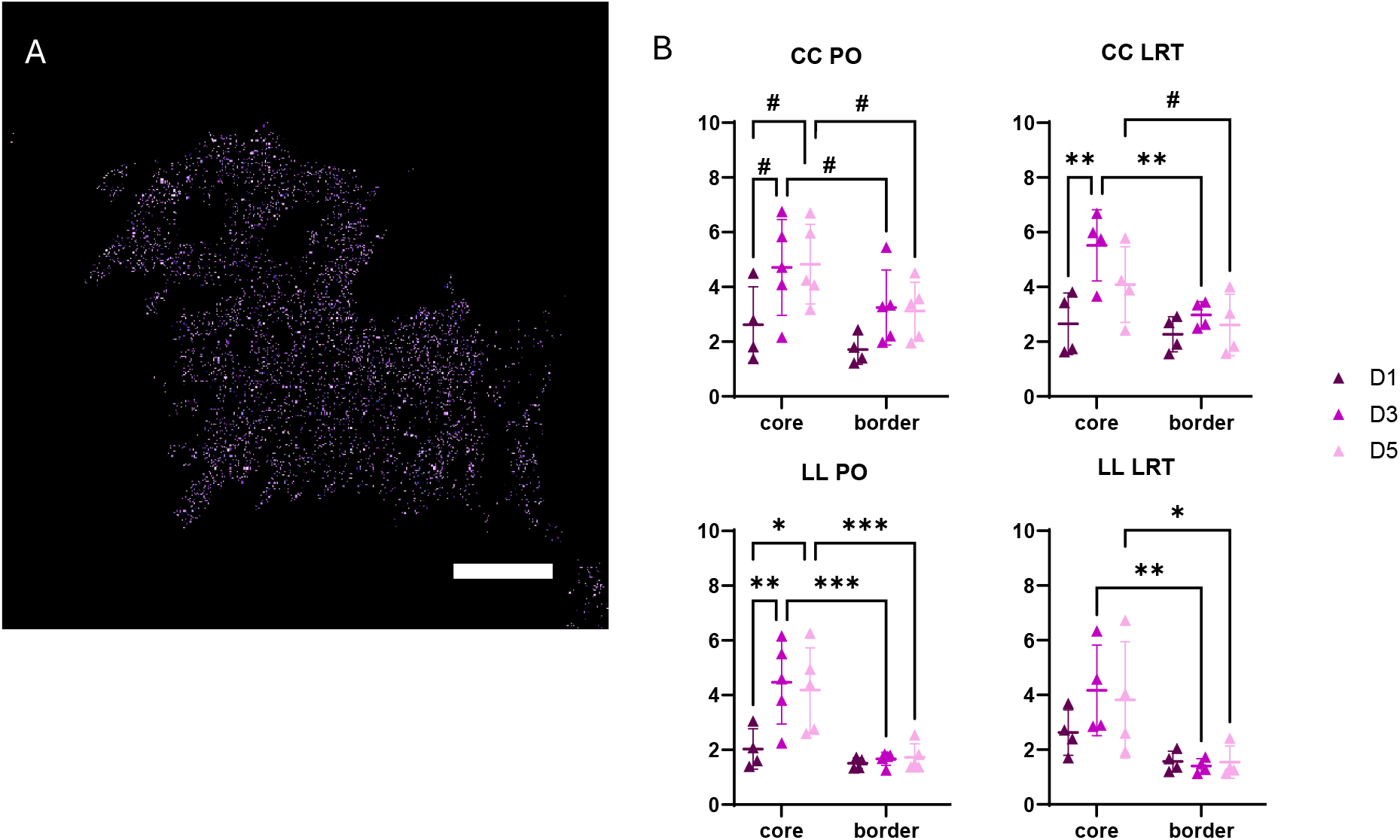
A) Cell nuclei for the H&E image of a representative day 3 permanent occlusion (PO) sample following color-based thresholding and isolation as well as the application of the core mask for the CC partitioning classification. Scale bar = 1mm B) Statistical analysis via 2-way ANOVA of cell area fraction between region (core and border were normalized to peripheral cell area fraction) and time point for the most conservative (CC) and liberal (LL) partitioning classifications. The LRT group included 4 hearts per day, and the PO group included 4 hearts at day 1 and 5 hearts at days 3 and 5. Pairwise comparisons via Tukey test: # p<0.1 ^*^ p <0.05, ^**^ p<0.01, ^***^p<0.001, ^****^ p<0.0001

Therefore, we conducted a 2-way ANOVA to assess the effects of region and time point (Figure 5B). The analysis indicated a significant difference in cell area fraction with both time point and region for both groups and all partitioning classifications. Specifically, p-values ranged from <0.0001 to 0.014 for region and from 0.013 to 0.023 for time point, except LC and LL for LRT group where time point was not significant. This analysis indicated a significant interaction between time point and region for the CL partitioning scheme for the late reperfusion samples only, p= 0.022. Paired comparisons via Tukey tests indicated the core cell area fraction was significantly greater than the border cell area fraction for both groups and all partitioning classifications for days 3 and 5, except CC PO where it neared significance at a p-value of 0.098 and 0.055 for days 3 and 5 respectively, and CC LRT day 5 where p=0.066. Significant temporal differences with region were only observed for the core. The core cell area fraction at day 1 was significantly or nearly significantly lower than that on days 3 and 5 for the PO samples in all partitioning classifications. In contrast, for the LRT samples the day 1 core area fraction was significantly lower than that on day 3 for all partitioning classifications (except LL), but only significantly or nearly significantly lower than that on day 5 for the CL and LC classifications (Figure S3). Notably, when the more conservative core definition was applied, core cell area fraction at day 5 trended downward in comparison to day 3 for the LRT group and remained elevated for the PO group. A Wilcoxon test indicated that normalized core and border cell area fractions were significantly larger than one for all partitioning classifications.

### 3.5 Collagen Area Fraction

For each sample in this study, we employed both PolScope and standard polarized imaging approaches to determine the collagen area fraction. Therefore, we overlaid regional masks to standardized polarized imaging of PSR stained samples to compute the amount of collagen in each sample region. A statistical analysis via 3-way ANOVA was conducted which revealed no significant 3-way interaction, but significant 2-way interactions were present between factors for all partitioning classifications. Consequently, we conducted 2-way ANOVAs to isolate and examine these interactions.

Statistical analysis via 2-way ANOVA between group and region indicated a significant difference in collagen density between the PO and LRT groups on days 1 and 3 and a significant difference in regional collagen density on days 3 and 5 regardless of partitioning classification (Figure 6A shows CC and LL; Figure S5 shows CL and LC). The interaction between group and region was not significant for any time point regardless of partitioning classification. At day 1, paired comparisons via Tukey tests indicated collagen density was significantly higher in the core of PO samples than LRT samples (Figure 6A and Figure S5; p-values ranged from 0.0059 to 0.017 depending on partitioning classification). This relationship reversed at day 3−collagen density in the core region of LRT samples was now higher than in PO samples and this reversal reached significance regardless of partitioning classification (p-values ranged from 0.0016 to 0.027). At days 3 and 5 collagen area fraction for the LRT group also exhibited regional differences. Core collagen density was higher than peripheral collagen density regardless of partitioning classification. This difference was statistically significant or nearly significant, with p-values ranging from 0.0069 to 0.071 for day 3 and from 0.0004 to 0.015 for day 5. Core collagen density was also greater than border collagen density for LRT samples at day 5, but this difference only reached significance for the CL and LL partitioning classifications. At day 5, the same regional pattern appeared in the PO group. Collagen density within the core was significantly greater than periphery and border for all partitioning classifications (p-values ranged from <0.0001 to 0.0007 for the periphery and 0.0048 to 0.027 for the border) except for the CC classification, where core collagen density was significantly higher than that in the periphery (p = 0.0017) but not the border (p = 0.67).

**Figure 6:**
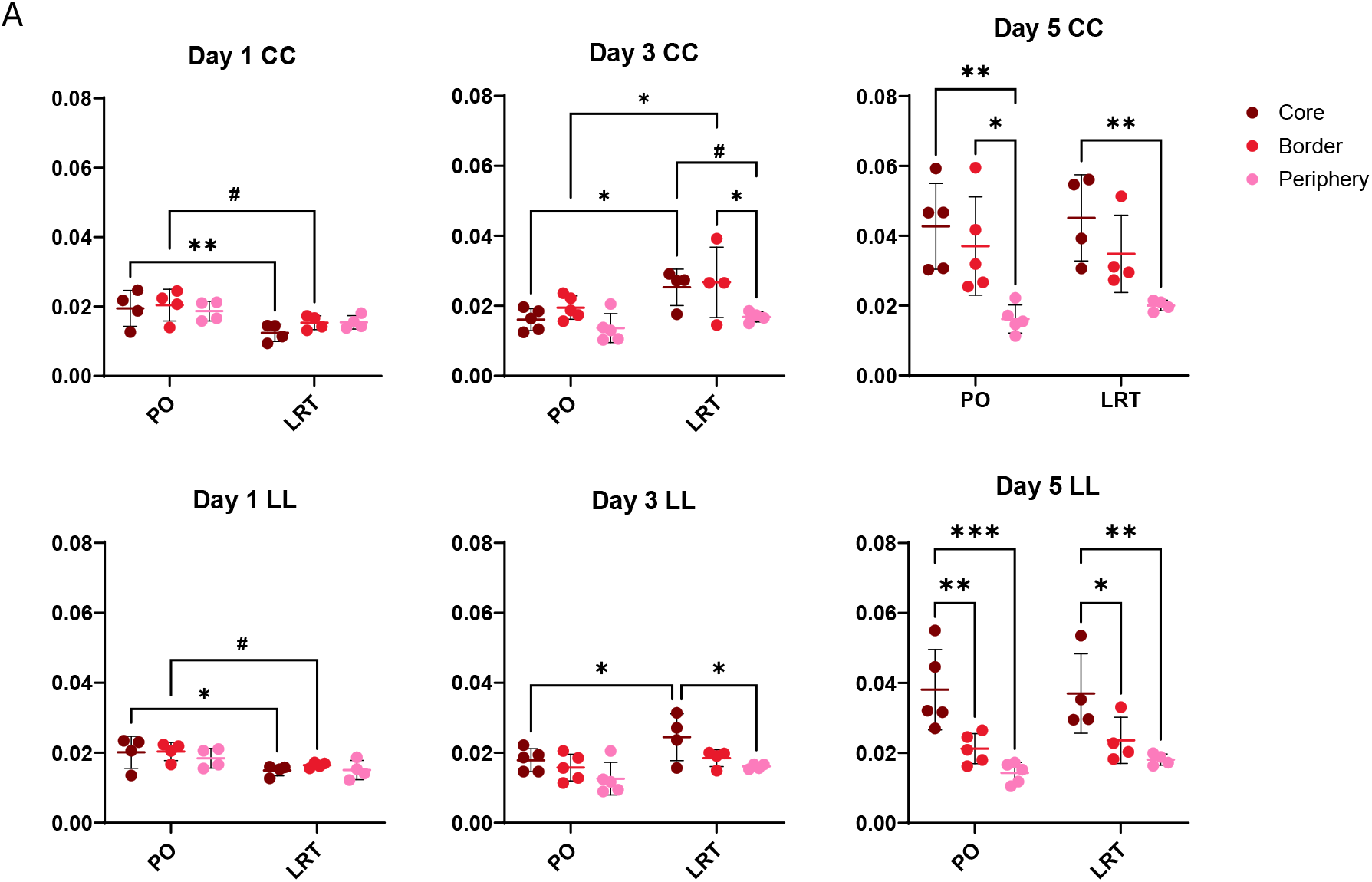
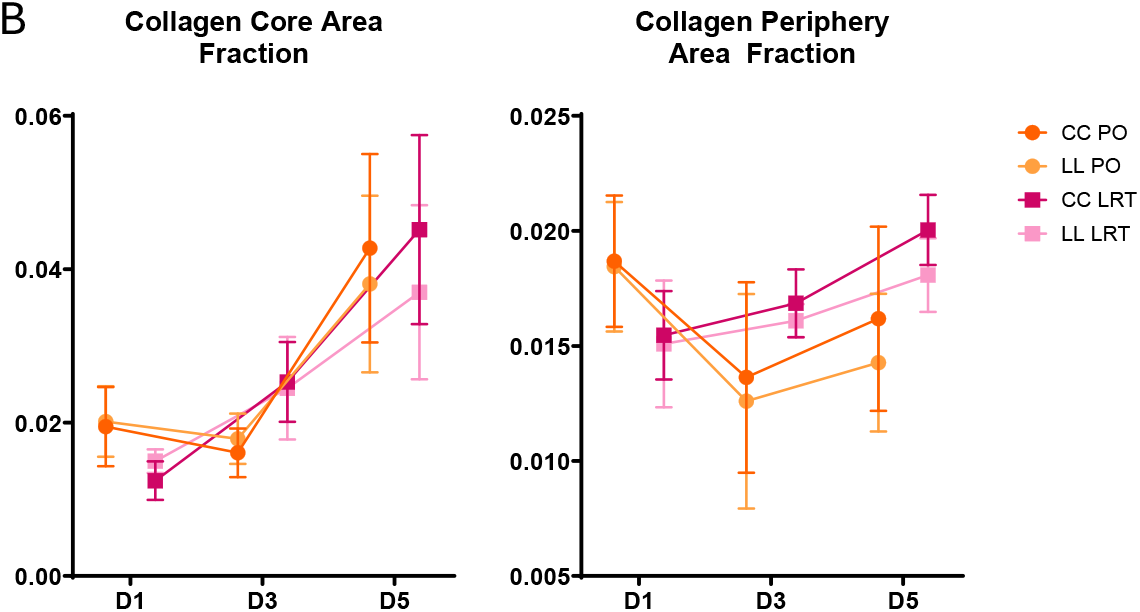
A) Statistical via a 2-way ANOVA of collagen density (as a percentage) between group (PO and LRT) and region (core, border, and global) for the most conservative (CC) and liberal (LL) partitioning classifications. There were significant differences between PO and LRT groups at days 1 and 3 and between regions at days 3 and 5. The LRT group included 4 hearts per day, and the PO group included 4 hearts at day 1 and 5 hearts at days 3 and 5. Pairwise comparisons via Tukey test: # p<0.1, ^*^ p<0.05, ^**^ p<0.01, ^***^ p<0.001, ^****^ p<0.0001. B) Core collagen density rose faster in LRT samples than PO samples (left, CC and LL partitioning classifications shown). Peripheral collagen density monotonically increased in LRT samples and was higher at days 3 and 5 than that of the PO samples (right). The LRT group included 4 hearts per day, and the PO group included 4 hearts at day 1 and 5 hearts at days 3 and 5. Pairwise comparisons via Tukey tests are detailed in Figure S6.

Statistical analysis via 2-way ANOVA between group and time point indicated no significant difference in collagen density between PO and LRT groups for any region regardless of partitioning classification (Figure S6). For the border region, collagen density differed significantly between time points, with p-values ranging from <0.0001 to 0.021 across partitioning classifications. For the periphery there was a significant interaction between group and time point, with p-values ranging from 0.036 to 0.050 across partitioning classifications. Core collagen density was significantly or nearly significantly higher at day 5 than at days 1 or 3 for both groups and all partitioning schemes according to pairwise comparisons via Tukey tests (Figure S6; p-values ranged from 0.0001 to 0.076 for days 3 and 5 and <0.0001 to 0.0060 for days 1 and 5). Figure 6B (left) emphasizes the different temporal trends between the groups – core collagen density remained low in the PO group for days 1 to 3 before increasing at day 5, whereas there was a steady increase in collagen density from day 1 to 3 and 3 to 5 in the LRT group. For both groups Tukey tests indicated core collagen density was significantly (p < 0.05) or nearly significantly higher (p < 0.10) higher at day 5 than days 1 or 3 for all partitioning classifications, but for the LRT group it was also significantly or nearly significantly higher at day 3 than day 1 for the CC and CL partitioning classifications. Notably, peripheral collagen density also differed temporally between groups (Figure 6B right). At day 1 peripheral density was significantly or nearly significantly higher than day 3 for the PO group (Figure S6; p-values ranged from to 0.022 to 0.053). Furthermore, at days 3 and 5 peripheral collagen density was higher in the LRT group than the PO, trending towards significance in most cases (p-values ranged from 0.096 to 0.13 for day 3 and 0.051 to 0.072 for day 5).

### 3.6 Collagen Color and Type in the Infarct Core

A shift in color in PSR stained images indicates changes in collagen type during infarct healing. The A value quantifies the PSR image coloration on a red-green scale, where collagen I is redder and collagen III greener. Figure S7 shows the A value for the core region of each sample at each time point. The 2-way ANOVA indicated no significant difference between the PO and LRT groups or between time points for either the conservative or liberal core definition. However, the temporal pattern in average A value for the PO and LRT groups differed. A decreased, indicating a shift from red (collagen I) on days 1 and 3 to green (collagen III) at day 5 in the PO group. In contrast, A decreased between days 1 and 3 followed by an increase between days 3 and 5 in the LRT group, suggesting a shift from red (collagen I) to green (collagen III) and back again.

## 4 Discussion

We conducted quantitative regional analysis to compare differences in the early structural remodeling of left ventricular tissue following permanent occlusion and restoration of blood flow following 3 hours of occlusion, simulating late reperfusion therapy. In general, late reperfusion therapy is associated with a reduction in incidents of rupture events post MI [14], [47], [60], [61]. However, the protective mechanism of this treatment is uncertain. Overall, our findings suggest that late reperfusion therapy results in accelerated post-MI healing. The infarct border width reduced more quickly and temporal changes in core collagen content and overall collagen coloring were consistent with faster healing. Therefore, late reperfusion therapy reduces the period of time where the LV wall is at risk of rupture events.

### 4.1 Partitioning Classification and Border Size

What constitutes the border zone around an infarct is frequently debated [47], [54], therefore we utilized four separate definitions ranging from the most conservative to most liberal to investigate the impact of these choices. As anticipated these definitions produced manyfold differences in both infarct size (Table S1) and border width (Figure 4A). While border width varied drastically between definitions, all exhibited a general temporal trend of decreasing border width for both permanent occlusion and late reperfusion samples. Additionally, at day 3, in most of the classifications, late reperfusion samples exhibited a smaller border width compared to permanent occlusion samples (Figure 4B). Studies investigating in-vivo rupture in mice reported ruptures occurred at the border and core, and ex-vivo mechanical testing of infarcted rabbit hearts reported an increased tendency to fail in the border and core [17], [38], [62]. Therefore, our observation that border size reduces more quickly in the late reperfusion group suggests a potential mechanism for the success of late reperfusion therapy in reducing rupture - the region most vulnerable to rupture decreases in size at a faster rate. An important distinction of our study was that infarct size and border width were quantified in a mid-wall slice along the longitudinal-circumferential plane of the LV (Figure 1), therefore transmural differences were not captured. Late reperfusion has not been shown to reduce infarct transmurality in rats [27] [60]. However, expansion is less clear, with one study reporting rats with late reperfusion exhibited thinning and dilation of the infarct region at two weeks [27] and another study reporting the scar segment length was shorter and thicker at 4 weeks [63].

### 4.2 Extracellular Matrix Changes

Out temporal observations in collagen density and type also support accelerated healing in the late reperfusion group via extracellular matrix remodeling. On day 1, the collagen density of the core and border regions was lower in the late reperfusion samples than the permanent occlusion samples (Figure 6B and Figure S5). This suggests the early remodeling during the necrotic phase of healing, which involves the breakdown of damaged myocardium, occurred more quickly in the late reperfusion samples [1], [2], [4]. Then, on day 3, the collagen density of the core region was higher in the late reperfusion samples than the permanent occlusion samples suggesting an earlier transition to the fibrotic phase of healing. Colormetric analysis of PSR stained samples further supports these observations. While not significant, there was a shift from red to green between days 1 and 3 in late reperfusion samples indicating collagen III deposition, whereas this shift occurred between days 3 and 5 for the permanent occlusion samples. Progression through the fibrotic phase of healing is characterized by an increase in the type I – to – type III collagen ratio [64]. Therefore, the shift in collagen coloration back towards red in the late reperfusion samples suggests that while collagen levels were not significantly different on day 5, some late reperfusion samples shifted towards collagen I deposition and progressed more quickly to the fibrotic phase of healing.

Towards the end of the necrotic phase of healing provisional extracellular matrix forms, characterized by the appearance of fibronectin, laminin, and collagen type IV, that is necessary for healing [1], [2], [4]. In our study, at day 1 non-collagenous fiber deposition was visible in PolScope images and present within the infarct core region of all samples. In the PO samples, however, this provisional matrix was localized to a very small area (Figure 7 right). Comparatively, the LRT samples exhibited a large diffuse network of small non-collagenous fibers Figure 7 (left). At day 3 PolScope images of the core regions of PO samples exhibited non-collagenous matrix at similar levels to day 1. In contrast, the provisional matrix was visible in only one LRT sample at day 3, which had a particularly large infarct core (6.6% or 17.5% area for conservative and liberal classifications, respectively). Therefore, beyond accelerating healing, late reperfusion therapy led to deposition of more non-collagenous matrix over a larger area during the necrotic phase of healing. Additional mechanical and structural analysis is necessary, however, to elucidate and quantify this difference in regional non-collagenous matrix deposition.

**Figure 7:**
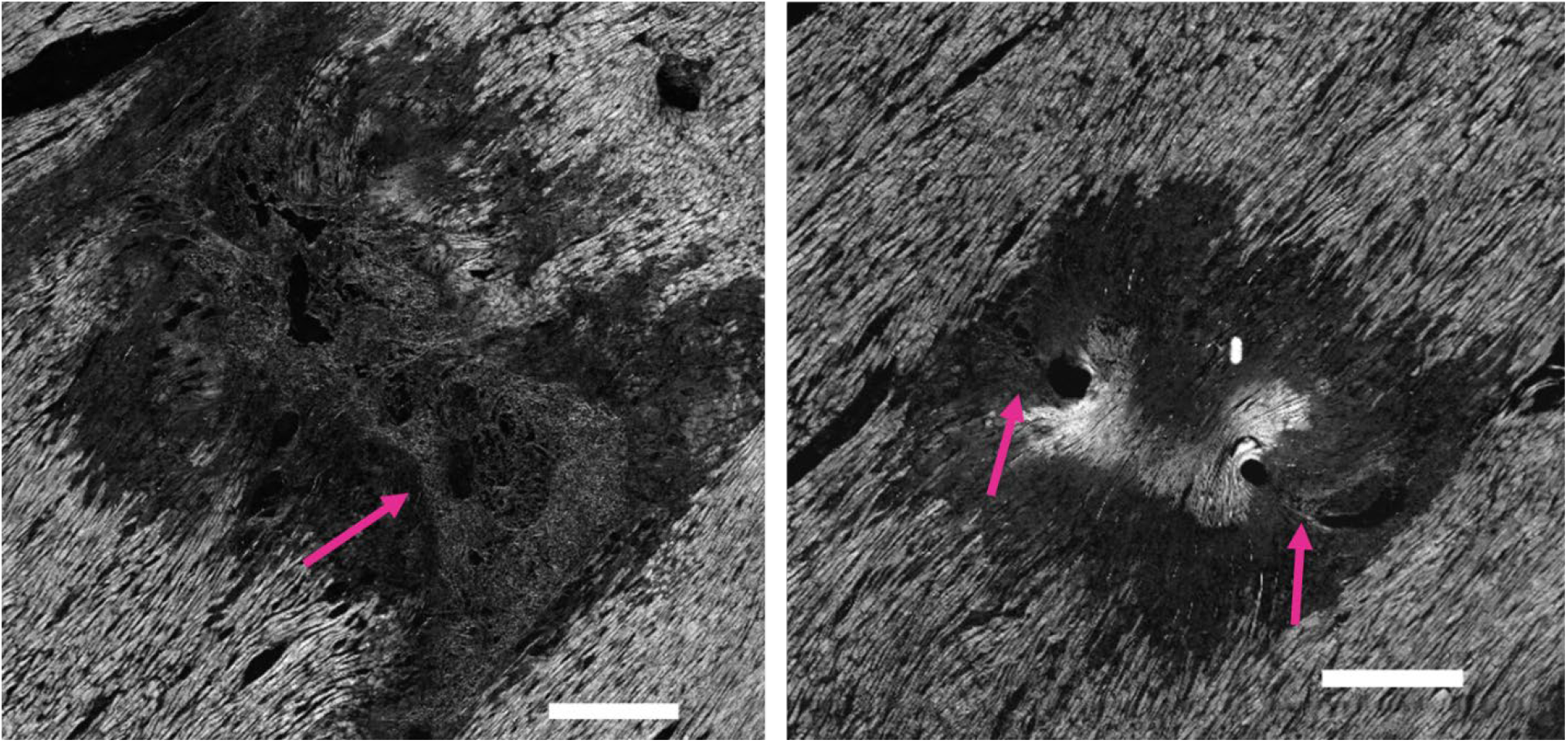
A representative PolScope image of an H&E stained section taken one day following late reperfusion (3 hours of ischemia, left) and permanent coronary artery occlusion (right). The day 1 reperfusion sample shows a large area of diffuse non-collagenous fibrous matrix (pink arrow), whereas the day 1 permanent occlusion sample shows limited non-collagenous fibrous matrix (pink arrows) localized near suture holes. Scale bar = 500μm

PolScope provided superior visualization of damaged tissue within the infarct core and border at day 1 (Figure 8A). Healthy cardiomyocytes were bright with clearly visible sarcomere units and intercalated discs, whereas damaged cells were darker and lacked these structural features. We initially conducted Masson’s trichrome staining to quantify infarct size, which has been widely used for this purpose [7], [46], [52], [63], [65], [66], [67], [68]. However, on day 1 changes in cardiomyocyte structure were insufficient in Masson’s trichrome stained samples to enable automated region detection (Figure 8B). Since damaged cardiomyocytes had not yet been removed and substantial changes in collagen structure were not yet present, PolScope was an ideal imaging method to quantify early ECM changes post-MI. By day 5, both tools produce similar results in delineating infarct region (Figure 9 C,D).

**Figure 8:**
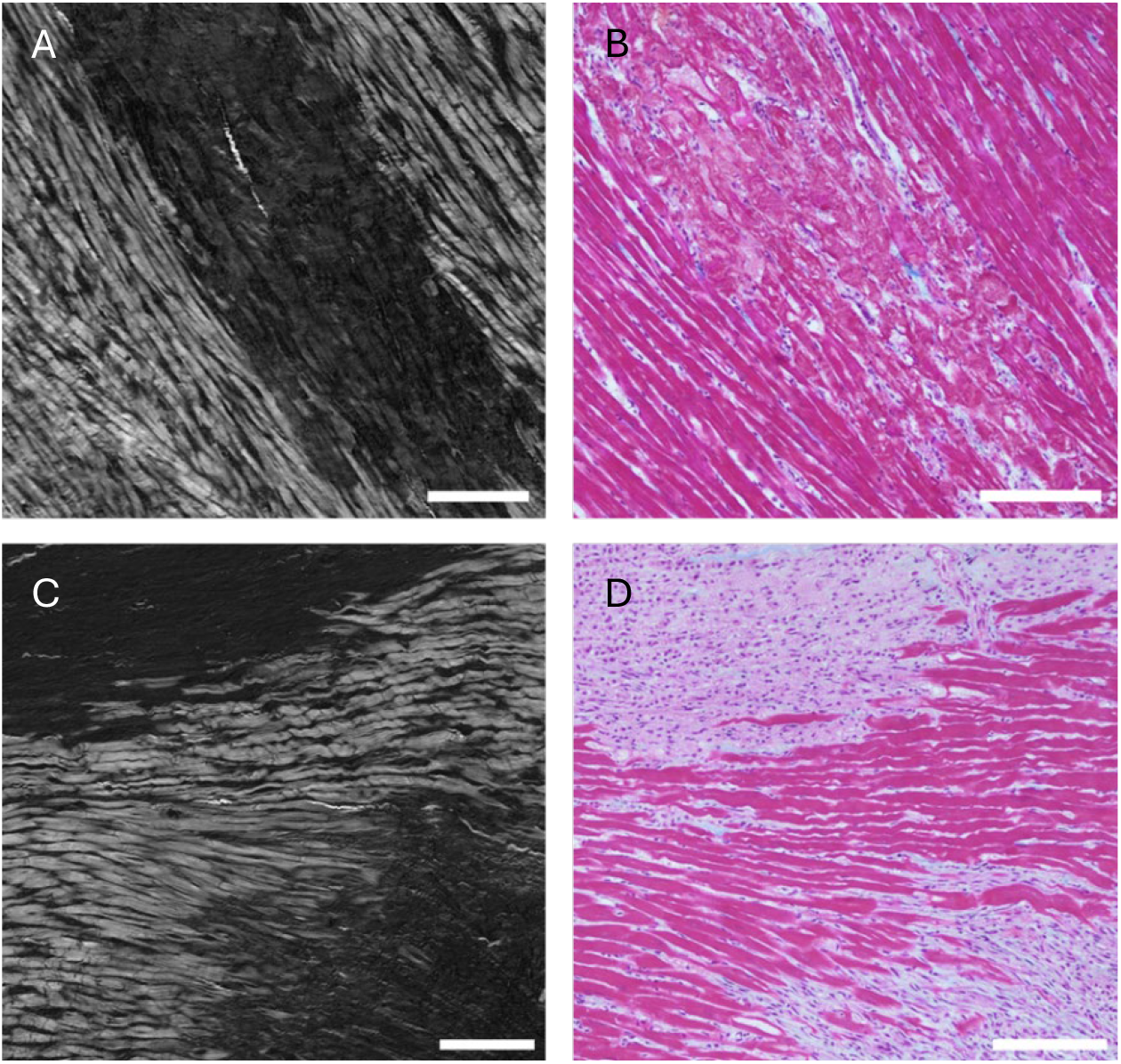
A, B) An infarcted region in a representative day 1 permanent occlusion sample. C, D) An infarcted region in a representative day 5 permanent occlusion sample. A, C) PolScope imaging of the H&E stained infarcted sample. B, D) Brightfield imaging of the trichrome stained infarcted sample. At day 1, damaged myocytes are clearly delineated in the PolScope image. While damage is visible in the brightfield trichrome image, the color variation needed for automated segmentation is not present. At day 5, both the PolScope and brightfield trichrome images clearly delineate the infarcted region. Scale bar = 150μm

### 4.3 Changes in Cell Density

The general trend of accelerated healing in the infarct with late reperfusion was less clear from the regional cell density results. In both sample groups for all partitioning classifications cellular density increased significantly from day 1 to day 3 (Figure S4), consistent with infiltration of immune cells and fibroblasts to facilitate remodeling during the necrotic and early fibrotic phases of healing [2], [4], [69]. For all partitioning classifications, core cell density remained elevated at day 5 for the permanent occlusion samples. However, when the conservative core definition was applied there was a statistical trend (p < 0.10) towards decreasing cell density in the core of late reperfusion samples, potentially supporting the acceleration of healing. Another novel result from our regional analysis was that the increase in cell density was concentrated in the infarct core. Only for the permanent occlusion samples and only in the most conservative of classifications did border cell density become significantly greater than the global cell density (Figure 5b). Therefore, our results do not support the hypothesis that late reperfusion somehow ‘spreads out’ the cellular response [70], [71]. Overall, our temporal results are consistent with previous studies on post MI healing which report increased cellular density over time [2], [4] but the spatial observations are novel.

Numerous cell types are involved in the infarct healing process, complicating regional differences in cellular infiltration and activity in a healing myocardial infarct irrespective of treatment. We observed no significant difference between the permanent occlusion and late reperfusion therapy groups at any single time point suggesting late reperfusion therapy does not affect overall cellular infiltration and density. However, it is possible late reperfusion therapy alters the infiltration rates of various cell types differently. Both macrophage and neutrophil upregulation have been reported to occur earlier in mouse models of late reperfusion therapy as compared to permanent occlusion following MI, but because they occur at different times the total cell content was unchanged [68], [70], [71]. A study in rats 4-days post-MI also reported a differential increase in cell type, with a higher levels of Ki-67+ cells in late reperfusion therapy samples and a higher level of TUNEL+ cells in permanent occlusion samples [63]. Additional studies including cell specific immunostaining could address this limitation. Lastly, low contrast between non-myocyte and cardiomyocyte coloration in H&E stained sections reduced our ability to automate raw cell density measurements. Therefore, we normalized cell density to the periphery to eliminate this effect.

## 5 Conclusion

In this study, we quantified the temporal changes in regional heterogeneity following permanent coronary occlusion and late reperfusion therapy. Using algorithmic image processing we were able to identify differences in cellular and collagen density between these groups both temporally and spatially. We found that late reperfusion therapy led to lower collagen density in the infarct core at day 1 and a higher collagen density in the core at day 3 compared to permanent occlusion. This temporal pattern indicates late reperfusion led to a more rapid progression through the necrotic phase of healing (when the infarct is vulnerable to rupture) and earlier progression to the fibrotic phase of healing (when collagenous matrix is deposited, and the infarct stabilizes). Colormetric analysis, while not significant, also suggested late reperfusion samples were shifting to collagen type I from type III at day 5 further supporting accelerated healing. Additionally, the border size decreased faster in late reperfusion therapy samples compared to permanent occlusion samples suggesting this accelerated healing also reduced the total tissue area most vulnerable to rupture. Lastly, we observed day 1 provisional non-collagenous matrix deposition over larger areas in late reperfusion therapy samples compared to permanent occlusion samples potentially suggesting an alternative mechanism to timing – more extracellular matrix material in the infarct core itself as another mechanism of rupture risk reduction. Together, these findings suggest that late reperfusion therapy may have multiple effects that reduce rupture risk including shortening the necrotic phase during which rupture is most likely, reducing the size of the infarct border more quickly, and potentially even producing provisional non-collagenous matrix over a larger area. Furthermore, the regional methodology we developed could be applied to discern the effects of more complex treatment plans including pharmacologic interventions and spatially variable approaches such as implantable cardiac patches or stem cell injections.

## Data availability

Data will be made available upon reasonable request.

## Declaration of Competing Interest

The authors declare no competing interests.

## Use of AI

The authors did not use generative AI or AI-assisted technologies in the development of this manuscript.

## Acknowledgements

This work was funded by a grant from the National Science Foundation Division of Civil, Mechanical and Manufacturing Innovation (ID, 2030173) to CMW and NIH U54CA268069 to KWE. The authors thank the University of Wisconsin Translational Research Initiatives in Pathology laboratory (TRIP) and the Experimental Animal Pathology Lab (EAPL), supported by the UW Department of Pathology and Laboratory Medicine, the University of Wisconsin Carbone Cancer Center (P30 CA014520), and the Office of The Director-NIH (S10 OD023526) for use of its facilities and services. The authors would also like to thank Daniel Pearce for his assistance with segmentation assignments, as well as Navya Sriram and Natalie Miller for their assistance imaging PSR stained slides.

## Notes

### Competing Interest Statement

The authors have declared no competing interest.

