## Supplement for "Regional and Temporal Changes in Early Structural Remodeling Following Myocardial Infarction via Semi-Automatic Image Analysis"

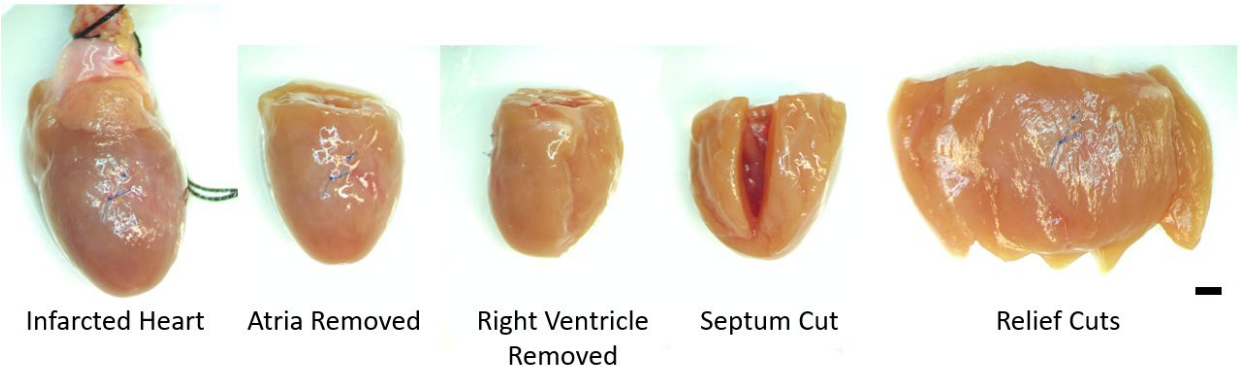


Figure S1: Following retrograde perfusion, the atria and right ventricle were removed from the LV. Then, the LV was cut open longitudinally along the septum and relief cuts along the apex were made such that the LV free wall could be flattened prior to histological sectioning. Scale bar = 2mm

Table S1: Percent area of the infarct core and border regions to the total sample area for all partitioning classifications from images of a mid-wall slice along the longitudinal-circumferential plane of the left ventricle wall. Entries are mean ± standard deviation.

| | CC | | CL | | LC | | LL | |
|---|---|---|---|---|---|---|---|---|
| | Core | Border | Core | Border | Core | Border | Core | Border |
| Day 1 | 4.63% ± 2.43% | 10.11% ± 1.75% | 4.63% ± 2.43% | 25.15% ± 4.58% | 14.74% ± 1.88% | 12.16% ± 3.42% | 14.74% ± 1.88% | 15.04% ± 4.06% |
| Day 1R | 5.04% ± 2.16% | 6.40% ± 2.12% | 5.04% ± 2.16% | 31.14% ± 7.69% | 15.06% ± 4.56% | 10.37% ± 2.19% | 15.06% ± 4.56% | 21.12% ± 7.34% |
| Day 3 | 5.37% ± 3.23% | 5.95% ± 2.11% | 5.37% ± 3.23% | 30.20% ± 10.46% | 14.53% ± 3.35% | 9.10% ± 1.96% | 14.53% ± 3.35% | 21.04% ± 10.63% |
| Day 3R | 4.61% ± 3.16% | 7.56% ± 2.40% | 4.61% ± 3.16% | 43.15% ± 16.41% | 17.65% ± 3.58% | 12.93% ± 2.65% | 17.65% ± 3.58% | 30.11% ± 15.32% |
| Day 5 | 3.14% ± 1.17% | 8.26% ± 2.01% | 3.14% ± 1.17% | 34.16% ± 18.42% | 12.66% ± 3.01% | 11.60% ± 4.06% | 12.66% ± 3.01% | 24.64% ± 16.26% |
| Day 5R | 6.85% ± 4.22% | 10.15% ± 4.49% | 6.85% ± 4.22% | 46.44% ± 9.28% | 18.82% ± 5.25% | 12.21% ± 4.28% | 18.82% ± 5.25% | 34.46% ± 6.79% |

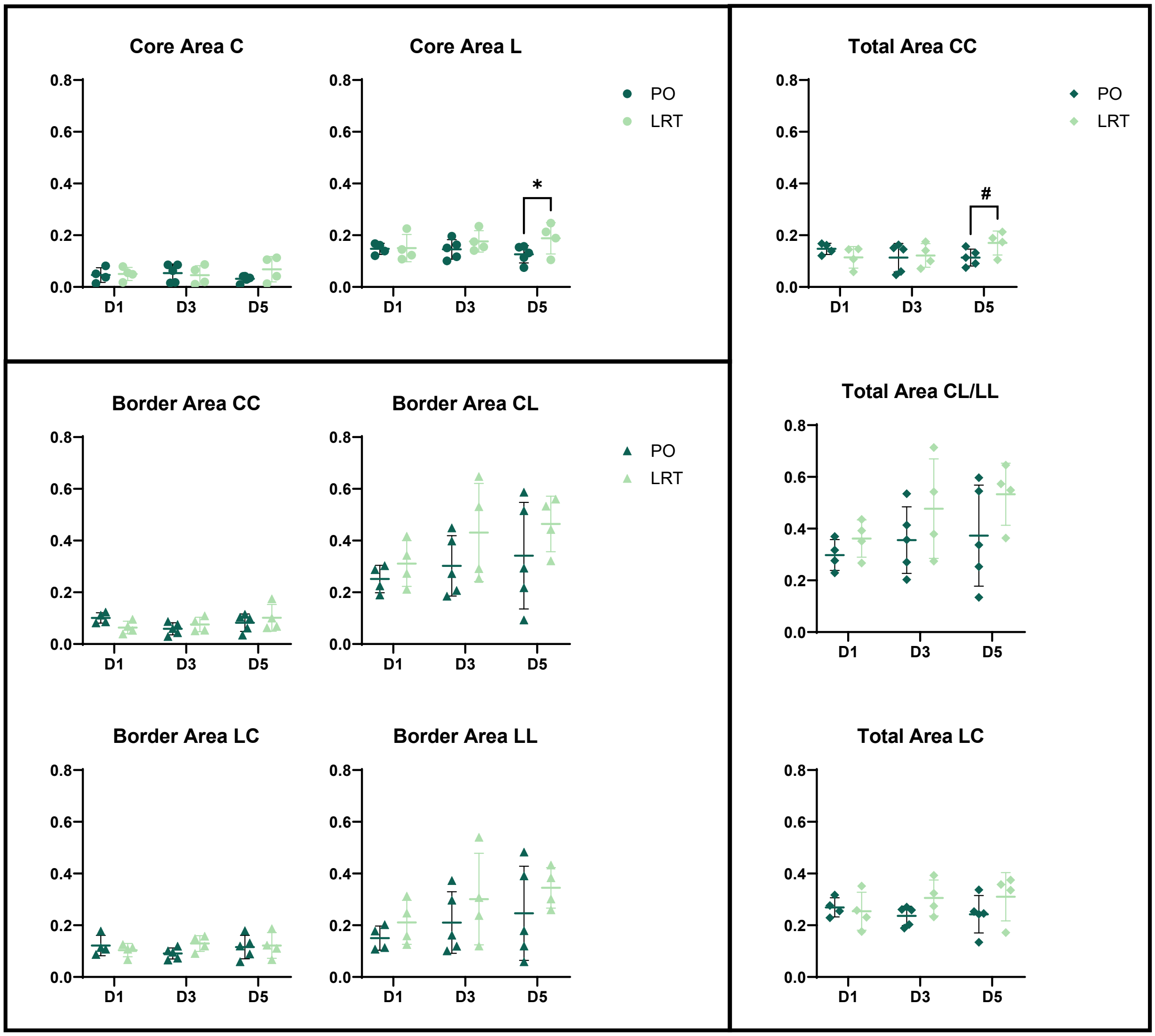


Figure S2: Core, Border and Total (Core + Border) Infarct areas as a percentage of the total area of the tissue on the stained slide. Statistical analysis via 2-way ANOVA indicated no significant differences in areas with group or time point . The LRT group included 4 hearts for each time point, and the PO group included 4 hearts at day 1, and 5 hearts at days 3 and 5 Pairwise comparisons via Tukey test: # $p<0.1$, * $p <0.05$

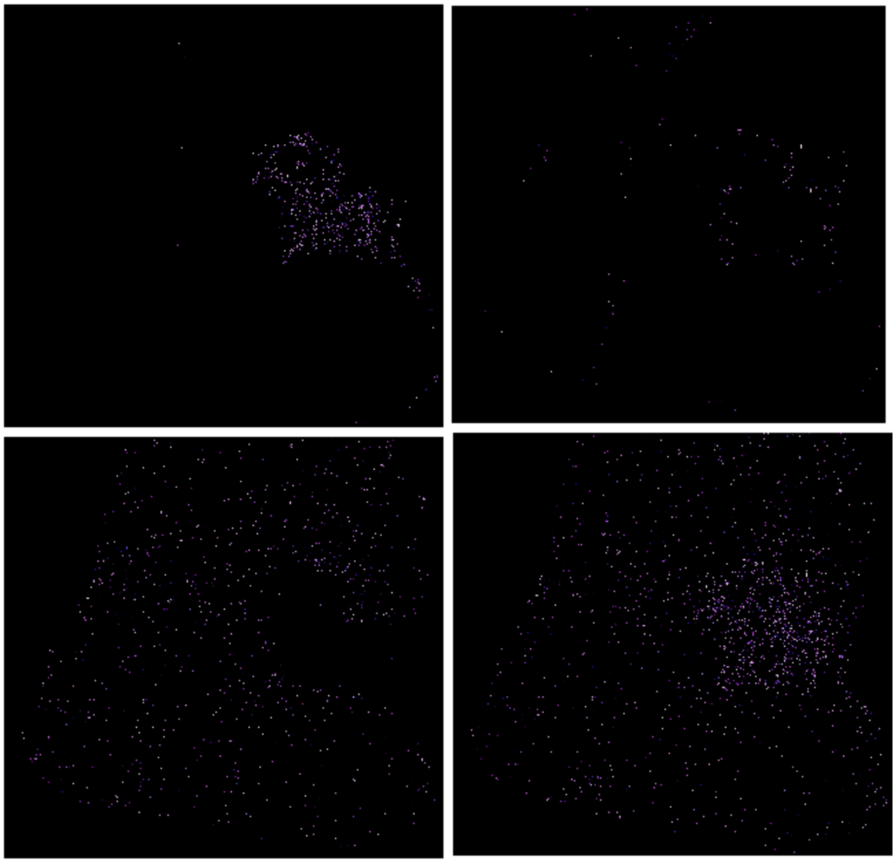

Figure S3: Cell nuclei for the H&E image of a representative day 3 PO sample following color-based thresholding and isolation as well as the application of the core mask (upper left), border mask (upper right), peripheral mask (lower left), or no mask (lower right) for the CC partitioning classification.

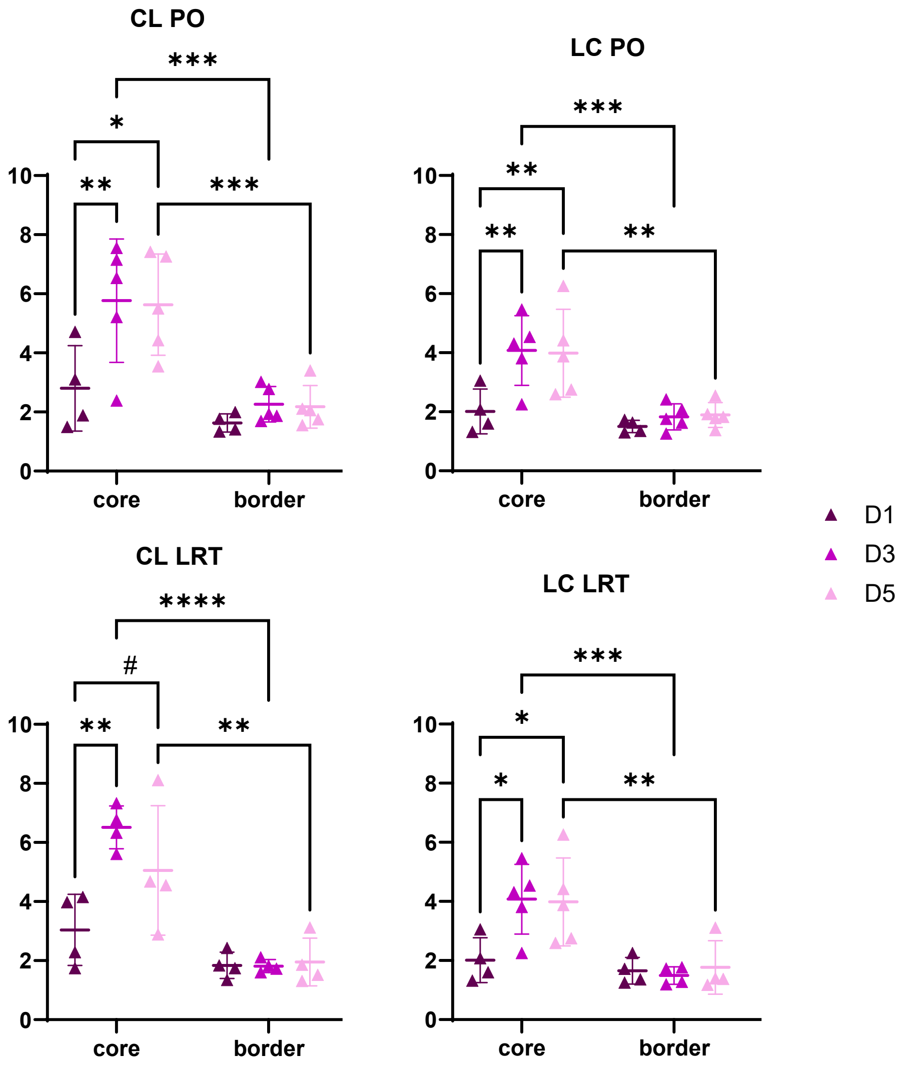


Figure S4: Statistical analysis via 2-way ANOVA of cell area fraction between region (core and border were normalized to peripheral cell area fraction) and time point for the CL and LC partitioning classifications. The LRT group included 4 hearts for each time point, and the PO group included 4 hearts at day 1, and 5 hearts at days 3 and 5. Pairwise comparisons via Tukey test: # p<0.1, * p <0.05, ** p<0.01, *** p<0.001, **** p<0.0001

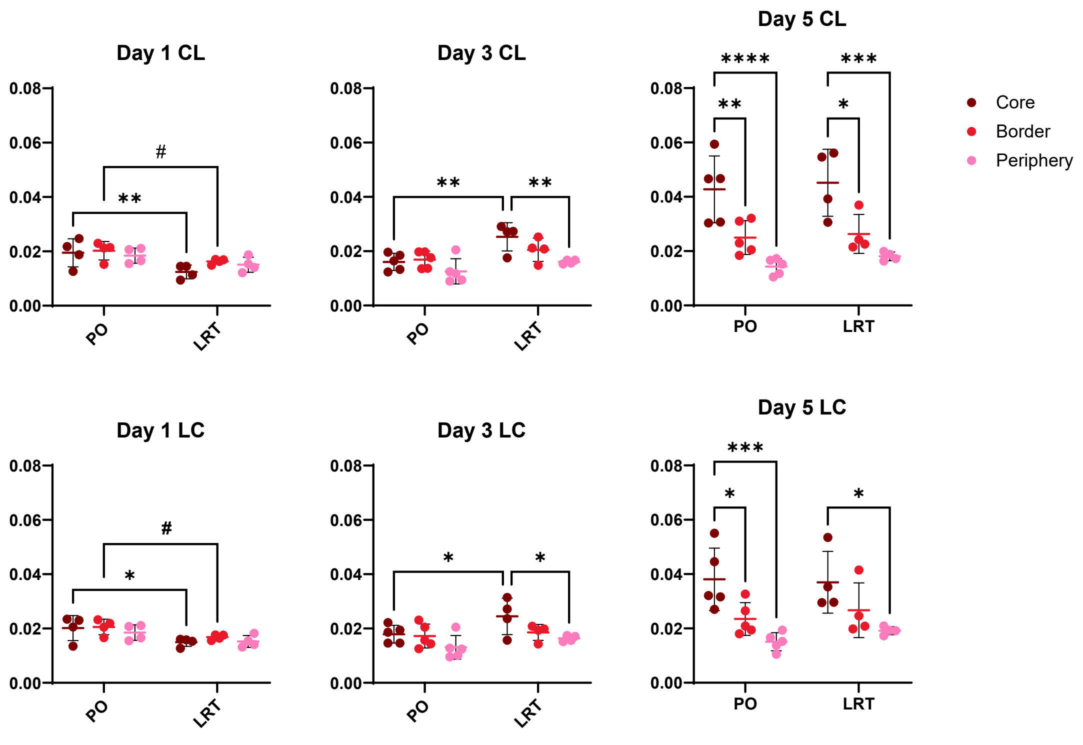


Figure S5: Statistical analysis via a 2-way ANOVA of collagen density (as a percentage) between group (PO and LRT) and region (core, border, and global) for the conservation core and liberal border (CL) and liberal core and conservative border (LC) partitioning classifications. There were significant differences between PO and LRT groups at days 1 and 3 and between regions at days 3 and 5. The LRT group included 4 hearts per day, and the PO group included 4 hearts at day 1, and 5 at days 3 and 5. Pairwise comparisons via Tukey test: # $p<0.1$, * $p<0.05$, ** $p<0.01$, *** $p<0.001$, **** $p<0.0001$.

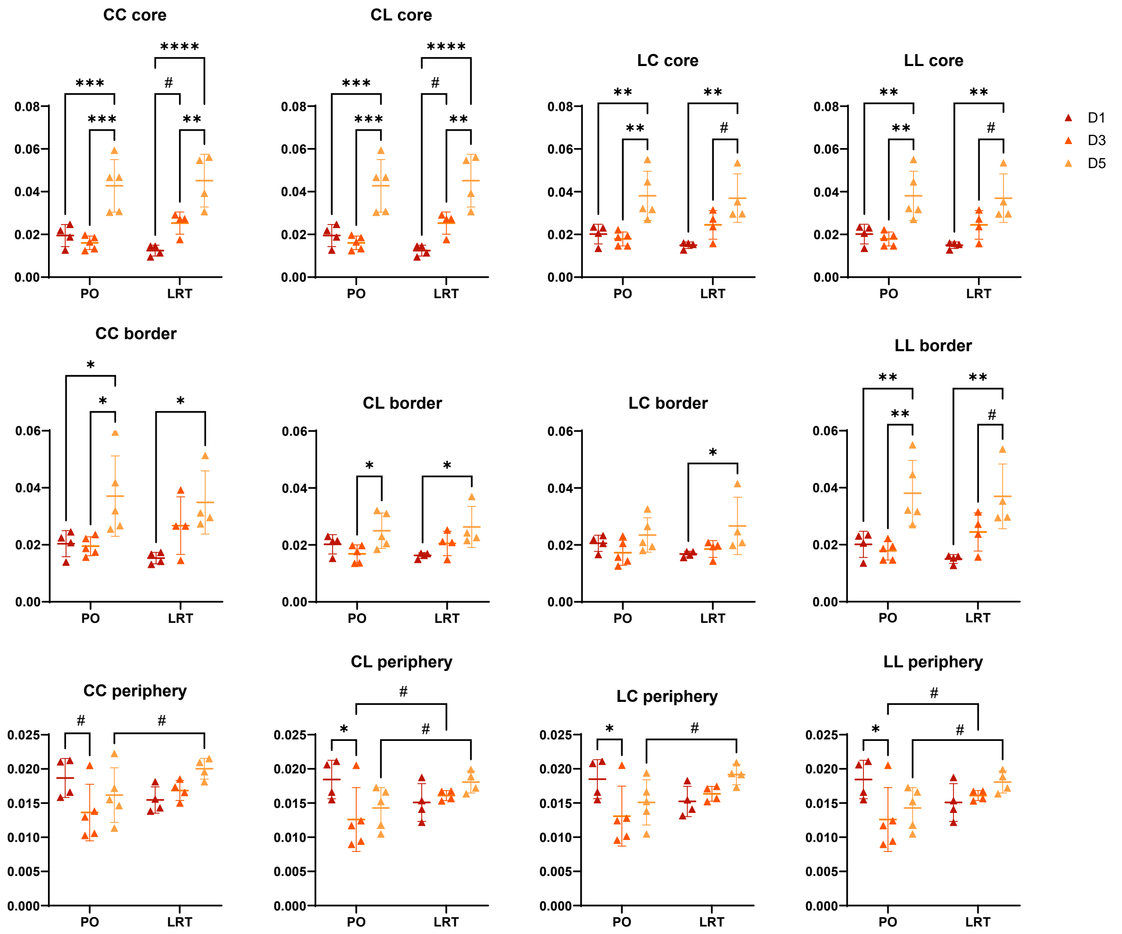


Figure S6: Statistical analysis via a 2-way ANOVA of collagen density (as a percentage) between group (PO and LRT) and time point (D1, D3, D5) for all partitioning classifications. The LRT group included 4 hearts per day, and the PO group included 4 hearts at day 1, and 5 hearts at days 3 and 5. Pairwise comparisons via Tukey test: # $p<0.1$, * $p<0.05$, ** $p<0.01$, *** $p<0.001$, **** $p<0.0001$.

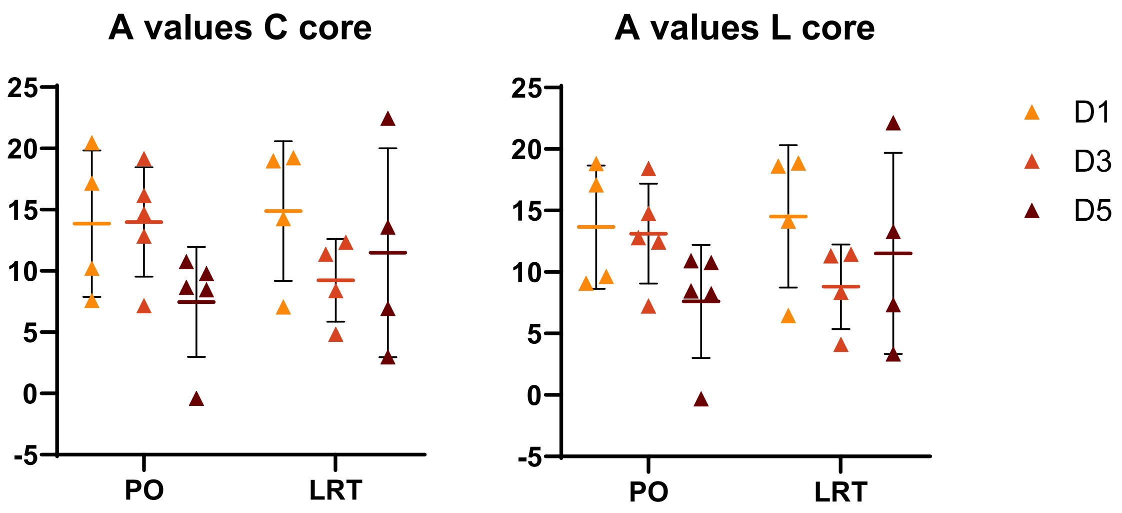


Figure S7: The shift in A value, i.e. red-to-green color transition, for conservative and liberal core partitioning classifications. the 2-way ANOVA indicated no significant difference between groups or time points, but the temporal trends in average A differed between the permanent occlusion (PO) and late reperfusion therapy (LRT) groups. The LRT group included 4 hearts per day, and the PO group included 4 hearts at day 1, and 5 hearts at days 3 and 5.
